# Selfish: Discovery of Differential Chromatin Interactions via a Self-Similarity Measure

**DOI:** 10.1101/540708

**Authors:** Abbas Roayaei Ardakany, Ferhat Ay, Stefano Lonardi

## Abstract

**Motivation:** High-throughput conformation capture experiments such as Hi-C provide genome-wide maps of chromatin interactions, enabling life scientists to investigate the role of the three-dimensional structure of genomes in gene regulation and other essential cellular functions. A fundamental problem in the analysis of Hi-C data is how to compare two contact maps derived from Hi-C experiments. Detecting similarities and differences between contact maps is critical in evaluating the reproducibility of replicate experiments and identifying differential genomic regions with biological significance. Due to the complexity of chromatin conformations and the presence of technology-driven and sequence-specific biases, the comparative analysis of Hi-C data is analytically and computationally challenging.

**Results:** We present a novel method called Selfish for the comparative analysis of Hi-C data that takes advantage of the structural self-similarity in contact maps. We define a novel self-similarity measure to design algorithms for (i) measuring reproducibility for Hi-C replicate experiments and (ii) finding differential chromatin interactions between two contact maps. Extensive experimental results on simulated and real data show that Selfish is more accurate and robust than state-of-the-art methods.

**Availability:** https://github.com/ucrbioinfo/Selfish

## 1 Introduction

Recent studies have revealed that genomic DNA in eukaryotes is not arbitrarily packed into the nucleus. The chromatin has a well organized and regulated structure in accordance to the stage of the cell cycle and environmental conditions (Ma *et al.*, 2015; Pederson, 1972). The chromatin structure in the nucleus plays a critical role in many essential cellular processes, including regulation of gene expression and DNA replication (Dixon *et al.*, 2012, 2015; Rao *et al.*, 2014; Sexton *et al.*, 2012).

Technological and scientific advancements in genome-wide DNA proximity ligation (Hi-C) have enabled life scientists to study how chromatin folding regulates cellular functions (Lieberman-Aiden *et al.*, 2009; Cavalli and Misteli, 2013; Chen *et al.*, 2015; Gorkin *et al.*, 2014). The analysis of Hi-C led to the discovery of new structural features of chromosomes such as topologically associating domains (TADs) (Dixon *et al.*, 2012; Zufferey *et al.*, 2018) and chromatin loops (Rao *et al.*, 2014; Cao *et al.*, 2018).

With the decreasing cost of Hi-C experiments and the higher availability of Hi-C data for different cell types in diverse conditions, there is a growing need for reliable and robust measures to systematically compare contact maps to discover similarities and differences. However, the comparative analysis of Hi-C data presents computational and analytical challenges which are due to technology-driven and sequence-specific biases. Technology-driven biases include sequencing depth, cross-linking conditions, circularization length, and restriction enzyme sites length (O’Sullivan *et al.*, 2013; Cournac *et al.*, 2012; Stansfield and Dozmorov, 2017). Sequence-specific biases include GC content of trimmed ligation junctions, sequence uniqueness, and nucleotide composition (Yaffe and Tanay, 2011). For instance, it is well-known that contact maps produced as a result of replicate experiments can contain significant difference solely due to these biases, which could be falsely interpreted as biological differences if these biases are not accounted for (Yardimci *et al.*, 2018). Several normalization methods have been developed to compensate for these biases and improve the reproducibility of Hi-C experiments, e.g., (Lieberman-Aiden *et al.*, 2009; Yaffe and Tanay, 2011; Knight and Ruiz, 2013; Imakaev *et al.*, 2012). While several computational methods have been proposed to extract statistically significant contacts from normalized contact maps (Rao *et al.*, 2014; Ay *et al.*, 2014; Cairns *et al.*, 2016; Ron *et al.*, 2017), their performance is still not entirely satisfactory due to the inherent complexity, inter-dependency and unaccounted biases in chromatin interaction data.

There are two major domains of application for the comparative analysis of Hi-C contact maps. The first application domain is focused on quantifying the reproducibility of Hi-C biological/technical replicate experiments (Yardimci *et al.*, 2018). For instance, Yang *et al.*, 2017 defined a reproducibility measure based on the *stratum-adjusted correlation coefficient statistic* defined on the unique spatial features of Hi-C data. Their method HiCRep (i) reduces the effect of noise and biases by applying a 2D averaging filter on the data, (ii) addresses the distance-dependence of Hi-C data by stratifying the data with respect to the genomic distance, (iii) calculates a Pearson correlation coefficient for each stratum, and (iv) aggregates the computed stratum-specific correlation coefficients using a weighted average.

In the same application domain aimed at quantifying reproducibility or concordance of contact maps, Ursu *et al.*, 2018 presented a method called GenomeDISCO to measure the differences between smoothed contact maps. GenomeDISCO represents contact map as a graph, where each node represents a genomic locus and each edge represents an interaction between two loci. Edges are weighted by the normalized frequency of the corresponding pairs of loci. GenomeDisco executes iteratively the two following steps: (i) processes the graph by random walks, which has the effect of denoising (smoothing) the data, (ii) computes the normalized difference between smoothed contact maps using the *L*_1_ distance between two contact maps.

The second application domain is aimed at finding statistically significant differences between contact maps for cells in different states (tissues, developmental states, healthy/diseased, time-points, etc). It is well-known that chromatin interactions that are mediated by specific protein can have distinct frequencies in different cell types or in different cell conditions (Patel *et al.*, 2012; Gong *et al.*, 2011). Differences in chromatin interactions can be associated with cell type-specific gene expression or mis-regulation of oncogenes or anti-oncogenes (Liu *et al.*, 2008; Schmitt *et al.*, 2016; Hnisz *et al.*, 2016).

Wang *et al.*, 2013 proposed the first method to discover differences in Hi-C contact maps. The authors used a simple fold-change of the normalized local interactions to discover that estrogen stimulation significantly impacts chromatin interactions in MCF7 cells. Building on this idea, Dixon *et al.*, 2015 proposed a method that (i) quantile-normalizes contact maps to compensate for the bias induced by different sequencing depth, and (ii) feeds the normalized differences of two contact maps (augmented by feature vectors representing epigenetic signals) to a Random Forest Model. Their method can (i) determine whether the epigenetic signal is predictive of changes in interaction frequency and (ii) discover which epigenetic signals are most predictive of changes in higher-order chromatin structure.

Stansfield and Dozmorov 2017 developed a non-parametric method to account for between-datasets biases. They used locally weighted polynomial regression to fit a simple model trained on the difference between the two datasets. Based on the assumption that the majority of the interactions should be relatively unchanged among similar Hi-C datasets and by centering the average difference to zero, loci which are far from the average are considered potential differential interaction.

Unlike other methods which assume independence among pairwise interactions, which holds true only for low resolution Hi-C data, Djekidel *et al.*, 2018 presented a method that takes into account the dependency of adjacent loci in higher resolutions. Based on the fact that interacting neighboring loci are known to inter-dependent, any structural difference can be detected by observing the differences in a neighborhood of the corresponding loci pair. In contrast, random noises tend to affect singular pairwise interactions only. By considering a three-dimensional space in which the *x* and *y* are the coordinates of the genomic loci and *z* is their pairwise interaction frequency, the authors define a chromatin interaction between two conditions to be *differential* when the intensity of the majority of *k*-nearest neighbors of (*x, y*) show a significant change.

In this work, we present new comparative methods for the analysis of Hi-C data based on the notion of self-similarity (Shechtman and Irani, 2007). We show that our self-similarity measure is robust to biases and does not need complex and computationally intensive normalization steps, such as MA (Dudoit *et al.*, 2002) or MD (Stansfield and Dozmorov, 2017). In the first part of the paper we show that our self-similarity measure can be used in the first application domain described above, i.e., as a tool to quantify the reproducibility of Hi-C biological/technical replicate experiments. In the second part of this paper, we show that our measure can also be employed in the second application domain, i.e., for finding statistically significant differences between Hi-C contact maps. Our method is based on the observation that interactions that are not independent of each other can be represented by their respective neighboring interactions. In other words, any structural difference between two contact maps not only affects the interaction value between its corresponding loci but also affects that of neighboring loci.

## 2 Materials and methods

Although existing methods for comparing contact maps vary widely, they all share the assumption that there exists common underlying global structures between two similar contact maps. We believe that this approach is fundamentally flawed, because the inherent biases present in the Hi-C data are very hard to model and completely eliminate. Here we propose to use the intrinsic self-similarity structure in contact maps to avoid dealing with the modeling problem.

In the application domain of object detection in complex visual data, the notion of self-similarity was first introduced by Shechtman and Irani, 2007. The idea of self-similarity on images can be explained as follows. Shechtman and Irani, 2007 showed that given two images of a certain object, the most relevant correlations between them are not necessarily the raw values of pixels (or an underlying model describing those pixel values) but the internal organization of self-similarities of local regions at similar relative geometric positions. In fact, for two images of the same object, we expect the relation between these local self-similarities to be more preserved than the similarities between the images.

### 2.1 Self-similarity and reproducibility

As said, existing methods for measuring the reproducibility of Hi-C experiments compute correlations or distances between normalized interaction frequencies of loci pairs, which is error-prone due to technology-driven and sequence-specific biases. Here we show that this comparison can be done indirectly by using self-similarity.

When we compare two contact maps that are expected to be similar, e.g. for two technical replicates of the same biological experiment, we expect to have similar internal layout of interactions. More precisely, given two contact maps *A* and *B* for two replicates, if we observe more chromatin interactions in block *α* than in block *β* in contact map *A*, we expect to have more chromatin interactions in block *α* than block *β* in contact map *B* as well, for several local choices of *α, β*. In other words, to measure similarity we do not need to depend on the absolute number of interactions in each contact map, rather we can rely binary comparison between many local interactions. Here we claim that the Boolean vectors representing binary comparison between local interactions encode enough information to define a similarity measure that can be used to quantify reproducibility for contact maps.

Henceforth, a Hi-C contact map is a *N × N* matrix where entry (*i, j*) in the matrix denotes the frequency of interaction between locus (or *bin*) *i* and locus (or *bin*) *j* in the genome. First, we slide a square block of size *N/k × N/k* along the main diagonal of the contact map using a stride of *N/*2*k* so each pair of adjacent blocks overlap by half of their size.For each position of the sliding block, we compute the sum of interaction frequencies inside the block. We store these sums in vector *B*, which has 2*k* components. We then compare all (ineq) pairs of block sums, and set the matrix *C*(*s, t*) = **I**(*B*_*s*_*> B*_*t*_) for all choices of (*s, t*) *∈* {1,*…,*2*k*} *×* {1,*…,*2*k*}, where **I** is the indicator function.

We claim that the matrix *C* is a compact representation of the interaction distribution along the main diagonal of the contact maps, which is robust to noise and biases (thus does not require normalization) because it relies on comparing entities that belong to the same contact map, and not across maps. We compute the similarity *S*(*A, B*) between contact map *A* and *B* as follows

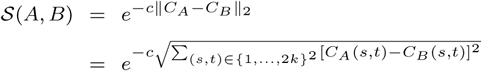

where *c* is a constant, *C*_*A*_ is the Boolean matrix for contact map *A, C_B_* is is the Boolean matrix for contact map *B*. The value of *k* should be chosen so that the size of the resulting blocks *N/k* is sufficient large to enclose important chromatin structures (e.g., TADs). Parameters *c* and *k* are determined experimentally.

### 2.2 Self-similarity and differential chromatin interactions

For the accurate detection of differential chromatin interactions (DCI), one needs to be able to distinguish true differences (which might have biological relevance) from differences caused by biases or other artifacts in the data. Since there is no ground truth for DCIs between cell types, conditions or developmental stages, there is no possibility of learning from real examples. The only differences that can be trusted are those that significantly exceed the differences observed between biological replicates. For this reason, we can also employ our self-similarity metric in a method for finding DCIs between two Hi-C contact maps. In our self-similarity representation described below, each interaction frequency is represented by a series of comparison between its surrounding local regions.

We first observe that DCIs have *locality* properties. If contact map *A* and *B* have a DCI at coordinate (*i, j*), this is not only reflected in the interaction difference of *A*(*i, j*) *-B*(*i, j*) but also in the neighborhood of (*i, j*). We call *impact region* the neighborhood affected by the DCI. We call *impact radius* the size of the neighborhood being affected, which is proportional to the magnitude of the DCI. We argue that what determines the statistical significance of a DCI in a particular location (*i, j*) does not only depend on the statistical significance of the difference *A*(*i, j*) *-B*(*i, j*) but also on the statistical significance of the difference between their region centered at (*i, j*). We argue that the most significant DCIs are not necessarily those locations that have the largest differences for a single interaction frequency. In fact, an isolated difference observed for a single coordinate (*i, j*) is more likely to be caused by an artifact in the data.

To incorporate locality information in our self-similarity representation, each interaction *A*(*i, j*) is represented by a linear combination of its neighboring interactions. To progressively penalize interactions which are farther and farther from (*i, j*), we weight these local interactions via a Gaussian filter c entered a t (*i, j)* (see Figure 1 for an example). By gradually increasing the size of the Gaussian filter, w e c apture impact regions with larger and larger radii. We denote with 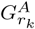 the matrix resulting from the convolution between a Gaussian filter w ith radius *r*_*k*_ and the contact map *A*. We first c ompute 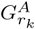 for a set o f *n* radii {*r*_1_, *r*_1_,…, *r*_*n*_}, and collect them in vector Γ*_A_* as follows

**Fig 1.**
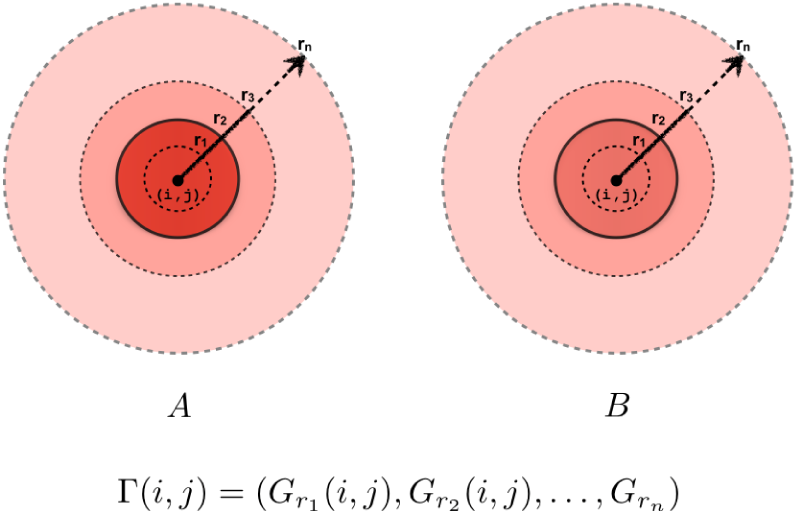
Our self-similar metric for representing chromatin interactions is obtained by first convoluting a contact map with a set of Gaussian filters with radii {*r*_1_, *r*_2_,…, *r*_*n*_}. The shade represents the intensity of the convolution for different radii. In this example, a sharp frequency change can be observed between radius *r*_*2*_ and *r*_*3*_ in contact map A but not in contact map *B* This difference can indicate a potential DCI.

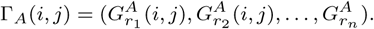

It is well-known that interactions in Hi-C contact maps are more frequent when the pairs of interacting loci are closer in genomic distance due to random polymer interactions driven by one-dimensional genome proximity. To account for this contact increase, we Z-normalize the interaction frequencies in *A* with respect to their genomic distances along each diagonal *d* as follows.

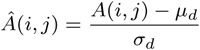

where *d* = *|j – i|*, and *µ*_*d*_, *σ*_*d*_ are the average and the standard deviation along the diagonal *d*, respectively.

If (*i, j*) is not a DCI between *A* and *B*, we expect vectors Γ _Â_(*i, j*) and 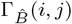 to exhibit similar trends along their components, because they represents aggregate interaction frequency in gradually increasing neighborhood centered at (*i, j*). If (*i, j*) is a DCI with impact radius *r*, we expect to observe a significant difference between the *k*-th Gaussian representations of that interaction, where *k* is the index of the radius *r*_*k*_ closest to *r*. Due to biases in the interaction frequencies across different contact maps, the difference between the two feature vectors Γ _*Â*_ and 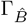 cannot be directly used to indicate the significance of a change. We address this issue by take advantage of self-similarity, i.e., by using local comparison of local regions in contact maps.

As it was done in Lowe 2004, instead of using Γ to define the behavior of interaction frequency across a set of impact regions, we use the first order derivative of Γ with respect to impact radius *r* to determine the behavior of interaction frequency, as follows.

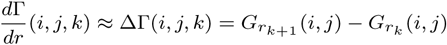

The difference 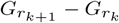 is sufficiently accurate to approximate the first order derivative 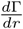 because the difference *r*_*k*+1_-*r*_*k*_ is small. By computing the first order derivative for various choices of the impact radii, we carry out a comparison of local contact map regions 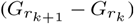.

Figure 2 shows the first order derivatives of Γ _*Â*_ and 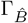 and the difference between them for the DCI reported later in Figure 7. Observe the sharp change between the derivatives at radius *r*_2_ which corresponds to a DCI with that radius.

**Fig 2.**
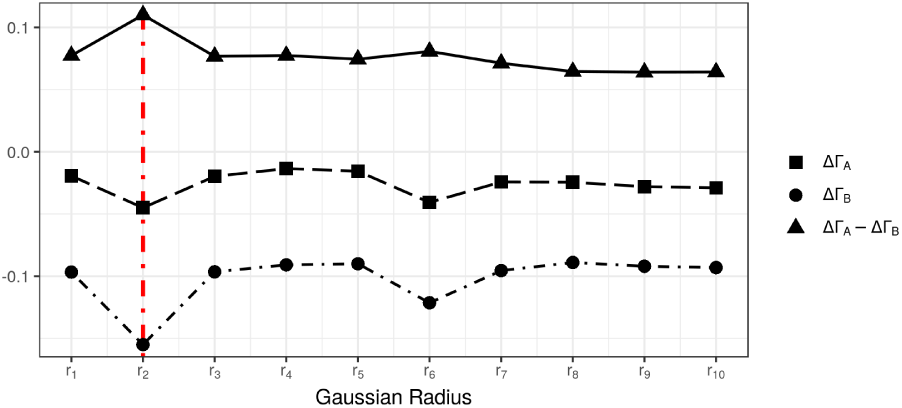
The first order derivatives of Γ*_A_*and Γ*_B_*and the difference between them. In this example, a large difference between ΔΓ*_A_*and ΔΓ*_B_*at radius *r*_2_ indicates a potential DCI.

In the last step of our algorithm, we find mean *µ* and standard deviation *σ* of a normal distribution fitted on the difference of the first order derivatives ΔΓ*_A_ -*ΔΓ*_B_* for each radius *r*_*k*_. Using these parameters of the normal distribution, we compute the *p*-value 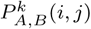 for location (*i, j*) and radius *r*_*k*_ as follows.

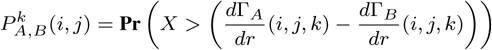

where *X* is normally distributed with parameters (*µ, σ*).

From the set of *k p*-values for each index (*i, j*), we choose the smallest as the *p*-value of the difference between two contact maps *A* and *B* at that index, as follows.

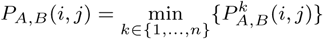

These *p*-values *P*_*A,B*_(*i, j*) are finally fed into the Benjamini-Hochberg algorithm to calculate the final probabilities (Benjamini and Hochberg, 1995).

## 3 Results

### 3.1 Reproducibility

We evaluated our reproducibility measure on a Hi-C dataset obtained from Schmitt *et al.*, 2016 that has a variable total number of interactions and resolution. The dataset consists of five different cell types hESC (H1), Mesendoderm (MES), Mesenchymal Stem Cell (MSC), Neural Progenitor Cell (NPC) and Trophoblast-like Cell (TRO). Each cell type each has two biological replicates. All experiments were carried out on a single chromosome (chromosome 1 for this work) with resolution 40 kb. The parameters *k, c* were experimentally set to *k* = 10 and *c* = 5. We compared our method SELFISH against the two state-of-the-art reproducibility methods in the literature, namely HiCRep (Yang *et al.*, 2017) and GenomeDISCO (Ursu *et al.*, 2018).

First, we assessed the effect of the total number of intra-chromosomal interactions captured by Hi-C experiment on different reproducibility measure. Given two biological replicates, we generated pseudo-replicates by first summing the two Hi-C matrices and then down-sampling the resulting matrix. Next, each individual replicate was down-sampled to a wide range of total interactions (10^5^, 5*×*10^5^, 10^6^, 2*×*10^6^, 5*×*10^6^, 10^7^). For each of these choice, we computed the reproducibilty score between all pairs.

Figure 3a-c illustrates the effect of the total number of interactions on the performance of reproducibility measures. A desirable feature for a reproducibility measure is to produce similar scores for the same pair of contact maps for different total number of interactions. Observe in Figure 3a-c that SELFISH is fairly invariant to the total number of interactions compared to HiCRep and GenomeDISCO, which failed to report stable reproducibility scores independently from the sequencing depth.

**Fig 3.**
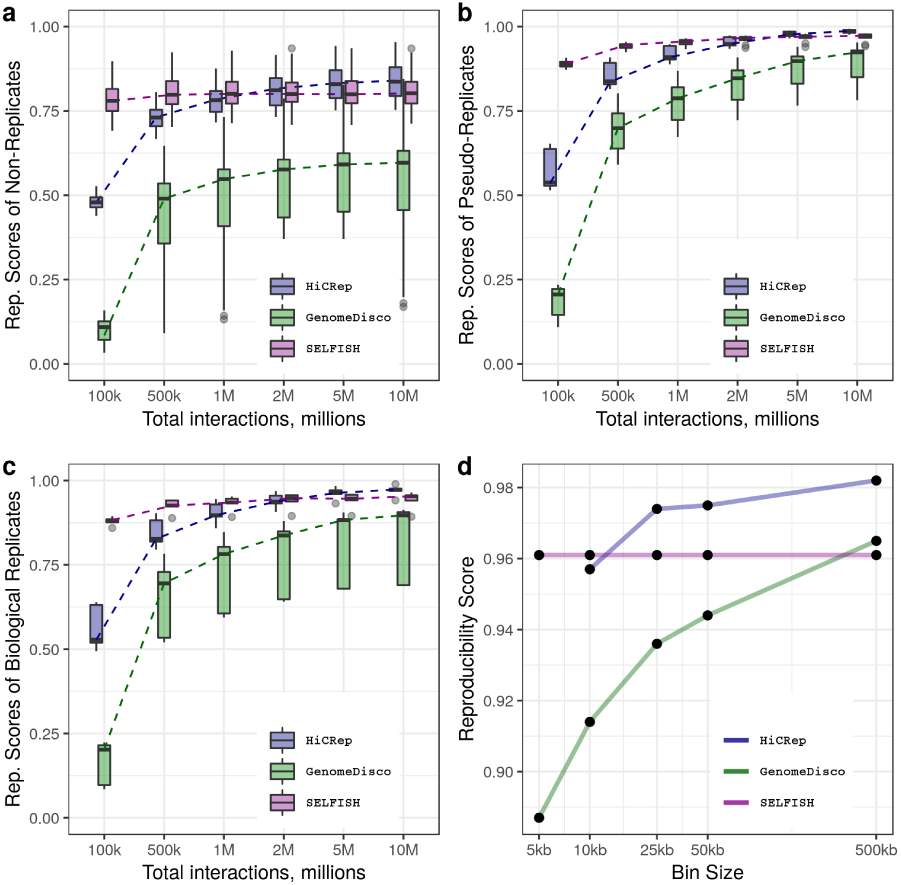
The effect of the total number of interactions on reproducibility score of (a) non-replicates, (b) pseudo-replicates and (c) biological replicates. (d) The effect of data resolution (bin-size) on reproducibility score of two replicates of cell type GM12878 from Rao et al. 2014

In the next experiment, we evaluated the effect of binning resolution of Hi-C data on the reproducibility methods. For this experiment, we used deeply sequenced Hi-C data of cell type GM12878 from Rao *et al.*, 2014. Again, a desirable property of a reproducibility score is to be robust to changes in resolution. Figure 3d shows that reproducibility scores for resolutions 5kb, 10kb, 25kb, 50kb and 500kb are very stable for SELFISH, whereas HiCRep and GenomeDISCO scores are resolution-dependent, in particular for GenomeDISCO.

These experimental results clearly indicate that SELFISH outperforms existing methods in terms of robustness to changes in sequencing depth and binning size. Both of these are very desirable features which can significantly simplify Hi-C data analysis in terms of quality control for reproducibility in replicate experiments.

We also compared the average running time of the three methods (see Table 1). The experiment is done on two replicates of GM12878 cell type from Rao *et al.*, 2014. The results show that SELFISH is by far more efficient than HiCRep and GenomeDISCO.

**Table 1.** Average running time of HiCRep, GenomeDisco and SELFISH for different choices of the data resolution

| Method | 500kb | 50kb | 25kb | 10kb | 5kb |
| --- | --- | --- | --- | --- | --- |
| HiCRep | 6s | 636s | 1045s | 34479s | * — |
| GenomeDisco | 7s | 2989 | 4933s | 30238s | 61004s |
| SELFISH | 0.75s | 85s | 184s | 345s | 474s |

\* HiCRep fails to run on 5kb data on a server machine with 256GB of RAM.

### 3.2 Differential Chromatin Interaction

We compared SELFISH to the current state-of-the-art method for detecting differential chromatin interaction called FIND Djekidel *et al.*, 2018. To the best of our knowledge, FIND is the only DCI detection method which works on high resolution Hi-C data by taking into account the chromatin interactions inter-dependency. Extensive experimental results in Djekidel *et al.*, 2018 show that FIND performs better than all prior methods for DCI detection. We experimentally set SELFISH’s parameters *n* = 10 and *r*^1^= 8.

We applied both methods on Hi-C contact maps for two cell types, namely GM12878 and K562, obtained from Rao *et al.*, 2014. We replicated some of the experiments in Djekidel *et al.*, 2018 in order to make a fair comparison. First, we analyzed the enrichment of epigenetic signals in the neighborhood of detected DCIs as well as the percentage of nearby genes having significant expression fold changes. We evaluated the enrichment of four well-known epigenetic markers, namely the binding of CTCF, POLII and P300, as well as the presence of histone modification H3K4me3. CTCF is widely accepted as a main driver of chromatin structure (Rao *et al.*, 2014; Tang *et al.*, 2015). We computed the enrichment of CTCF differential peaks (peaks that are different between two cell types) around detected DCIs. For this part of the analysis we directly obtained the detected DCIs for the mentioned dataset from Djekidel *et al.*, 2018.

To compute the enrichment of each marker in the neighborhood of the detected DCIs, we calculated the distance of marker peaks to their closest anchor of DCIs. Figure 4 shows the enrichment of epigenetic markers near DCIs, highlighting especially for CTCF and H3K4me3 that these marks are more enriched around the reported DCIs for SELFISH compared with those detected by FIND, even though the number of reported DCIs for SELFISH is twice as large (30456 vs. 14131).

**Fig 4.**
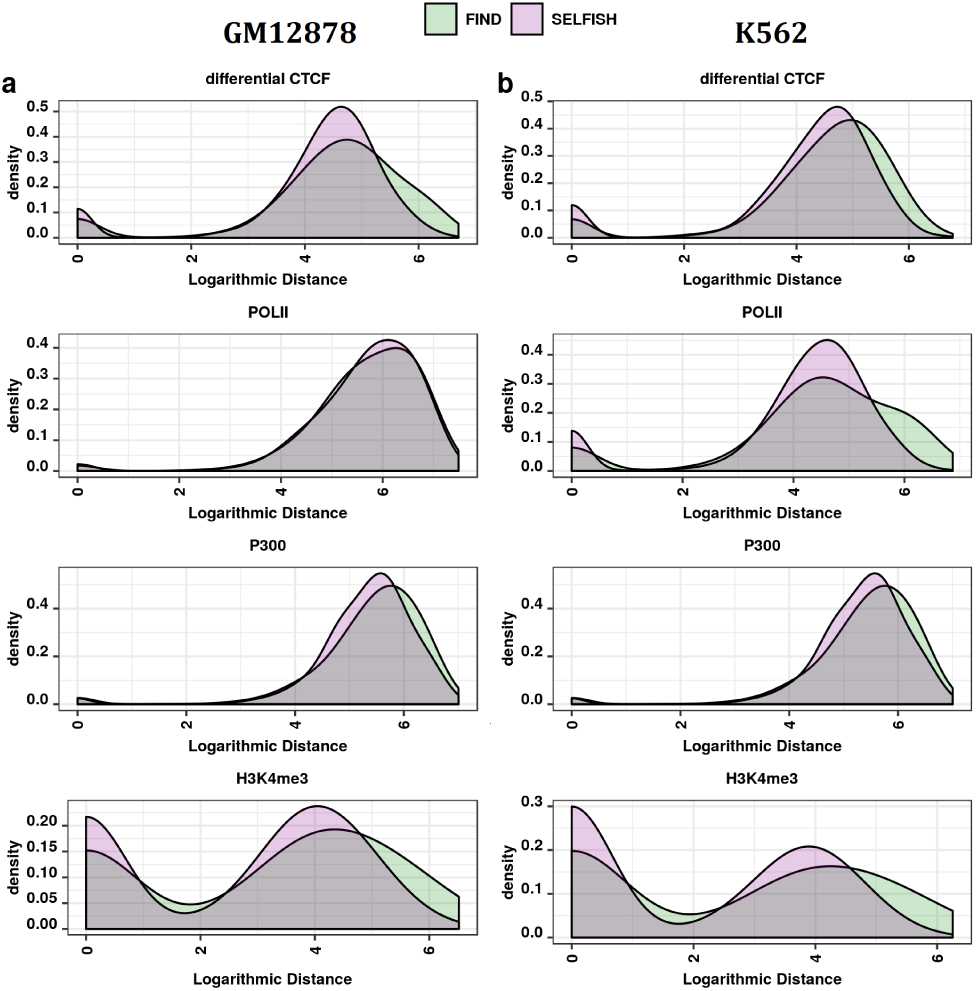
Enrichment of differential transcription factor binding and epigenetic marks (CTCF, POLII, P300 and H3K4me3) around reported DCIs for (a) cell type GM12878 and (b) cell type K562.

We also calculated the expression fold change of nearby genes for the two cell types. We first determined the set of genes which have overlap with any of the detected DCIs’ anchors. Then we computed the percentage of those genes having an expression fold change of two or greater. For the set of genes overlapping FIND’s DCIs, 71.46% of them were over-expressed. For SELFISH, 78.78%, were over-expressed. This analysis confirmed that the differences in chromatin structure are strongly associated with the changes in gene regulation. However, the DCIs detected by SELFISH have stronger associations to differences in gene regulation than FIND.

To quantify how well the Hi-C data supported the detected DCIs between two cell types GM12878 and K562, we generated a modified aggregate peak analysis (APA) plots (Rao *et al.*, 2014; Phanstiel *et al.*, 2015). The interaction frequencies in contact maps are first Z-normalized by diagonal as explained in Methods section. Then, for each detected DCI we calculated the interaction differences between the contact maps in a *±*50*kb* neighborhood. By averaging over all DCIs, we computed the APA plot for differences. The differential APA score, i.e. the value of the central index in the plot compared to neighboring regions, shows how different the interactions are at reported DCIs with respect to their expected interaction frequency with that genomic distance.

Figure 5 shows that SELFISH produces the expected Gaussian-shaped plot around its reported DCIs while the DCIs from FIND failed to generate a similar pattern. Correspondingly, the computed APA score is higher for SELFISH (4.46) compared to FIND (3.06) suggesting a stronger detection of DCIs. Finally the peak pixel of the APA plot for FIND (score of 4.17) is not centered on the called DCI pairs suggesting that SELFISH performs better at pinpointing anchor points of chromatin interactions at high resolution.

**Fig 5.**
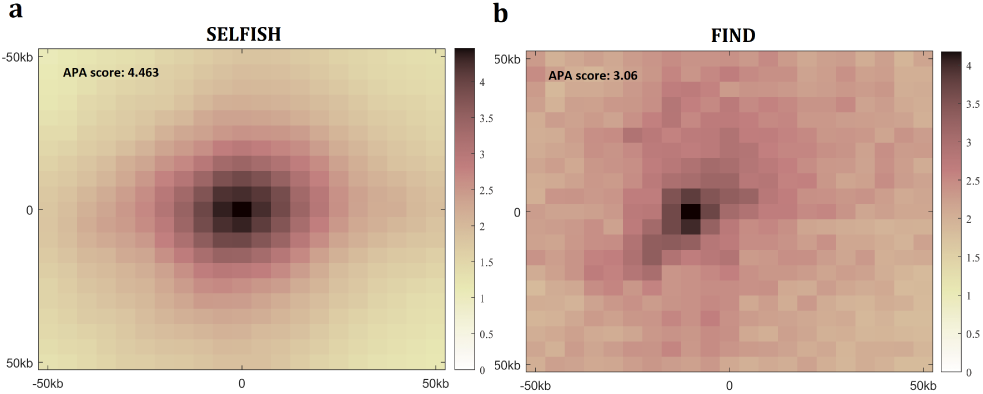
A modified APA plots for reported DCIs between two cell types GM12878 and K562 by (a) SELFISH and (b) FIND.

To further investigate the accuracy of detected DCIs we used simulated Hi-C data generated by the method proposed in Zhou *et al.*, 2014. We generated a set of 100 pairs of simulated contact maps, each of which had known location for DCIs. After running SELFISH and FIND on these simulated datasets, we obtained a *p*-value for each DCI location for all 100 simulated pairs of contact maps. Given the *p*-values and the true locations of DCIs (true positives) we computed a precision-recall curve for each simulated pair of contact maps. We used the threshold averaging method proposed by Fawcett, 2006 to combine the 100 precision-recall curves to get the overall performance curve. We thresholded over the ratio of all indices in the contact map used for computing the precision and recall for each simulated pair. To combine the curves, we averaged all 100 calculated precision and recall values for each threshold. Figure 6a-c shows the performance of both methods for 2-fold, 5-fold and 10-fold DCIs. The vertical and horizontal bars represent the 95% confidence interval for precision and recall at that threshold respectively. For all fold change settings, SELFISH performs better than FIND confirming its superior performance on real Hi-C datasets. The performance difference is most striking for small fold change values, which are more relevant for comparisons of real Hi-C datasets and yet have very large effect on gene regulation (Greenwald *et al.*, 2018). It is also important to note that SELFISH’s performance is quite consistent across different samples as indicated by small confidence intervals.

**Fig 6.**
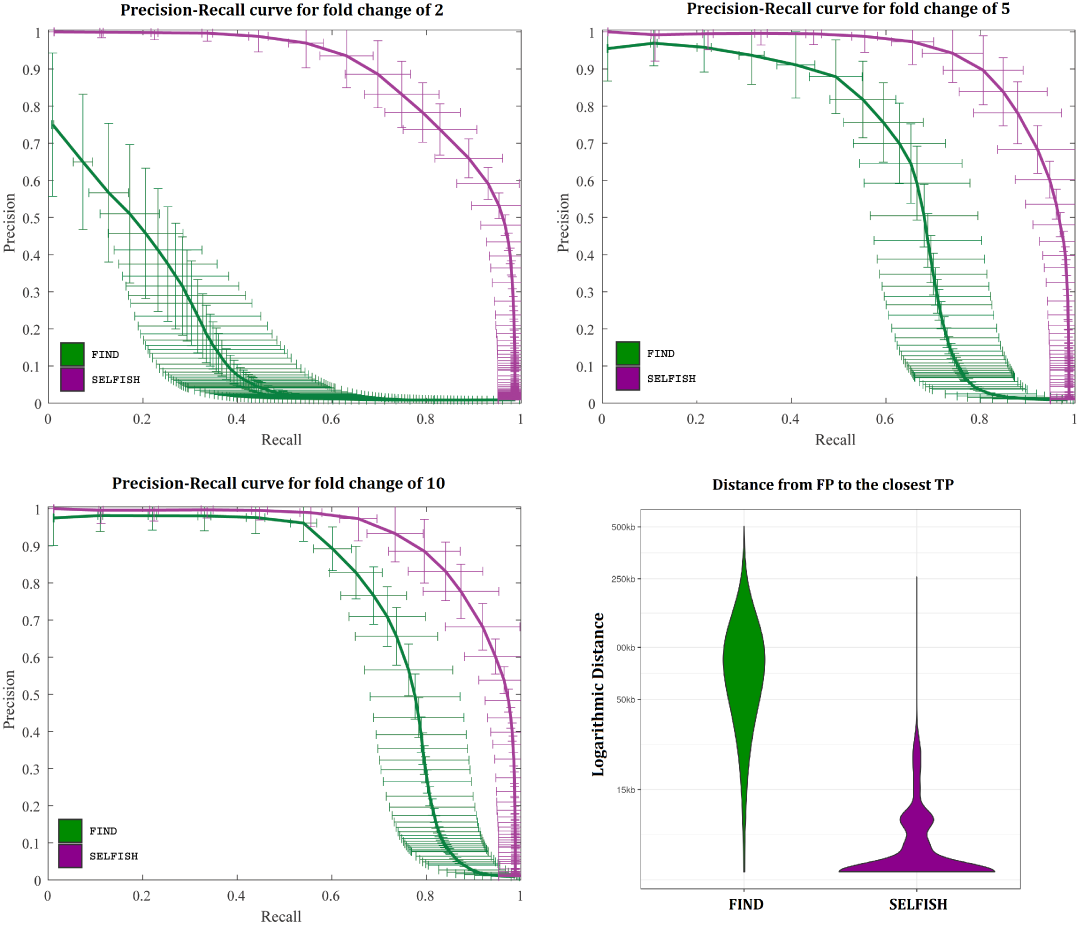
Precision-Recall curves for SELFISH (magenta) and FIND (green) for (a) 2-fold, (b) 5-fold and (b) 10-fold DCIs. The vertical and horizontal bars represent the 95% confidence interval for precision and recall at that threshold respectively. (d) The distribution of distances of FPs to closest TPs.

Figure 6d shows the distribution of distances between false positive DCIs produced by SELFISH and FIND to true DCIs (true positives). To generate this figure we set the number of returned DCIs equal to the number of true DCIs. Then, for each falsely detected DCI we calculated its distance to the closest true DCI. These results clearly show that most false positives for SELFISH are located in close proximity of true DCIs confirming their relevance to true differences and their non-random distribution. On the other hand, false positive DCIs detected by FIND are farther from true DCIs and are more scattered in the contact map.

In our final experiment to assess the performance of two methods, we tested SELFISH and FIND on a real test case from Bonev *et al.*, 2017. Figure 7 shows a 2-Mb region around the *Brn2* promoter (also known as *Pou3f2*) for mouse embryonic stem cells (ES) and neuronal progenitor cells (NPC). Dashed circles show the contact between the *Brn2* promoter and an NPC specific enhancer. Insets show the magnified view of this contact. Observe that the contact between the promoter and enhancer is strongly present in the NPC cell (Figure 7a) in contrast with the ES cell in which this interaction is weak (Figure 7b). The mentioned contrast shows itself as a subtle but important difference of interactions between two cell types. The highlighted regions of the epigenetic signals show the difference in the specified regions between two cell types. Detected DCIs by SELFISH and FIND for *q*-value *<* 10*-*^4^ are shown in magenta and green squares respectively. Observe that SELFISH can identify this contact region as a DCI between two cell types, but FIND fails to detect it.

**Fig 7.**
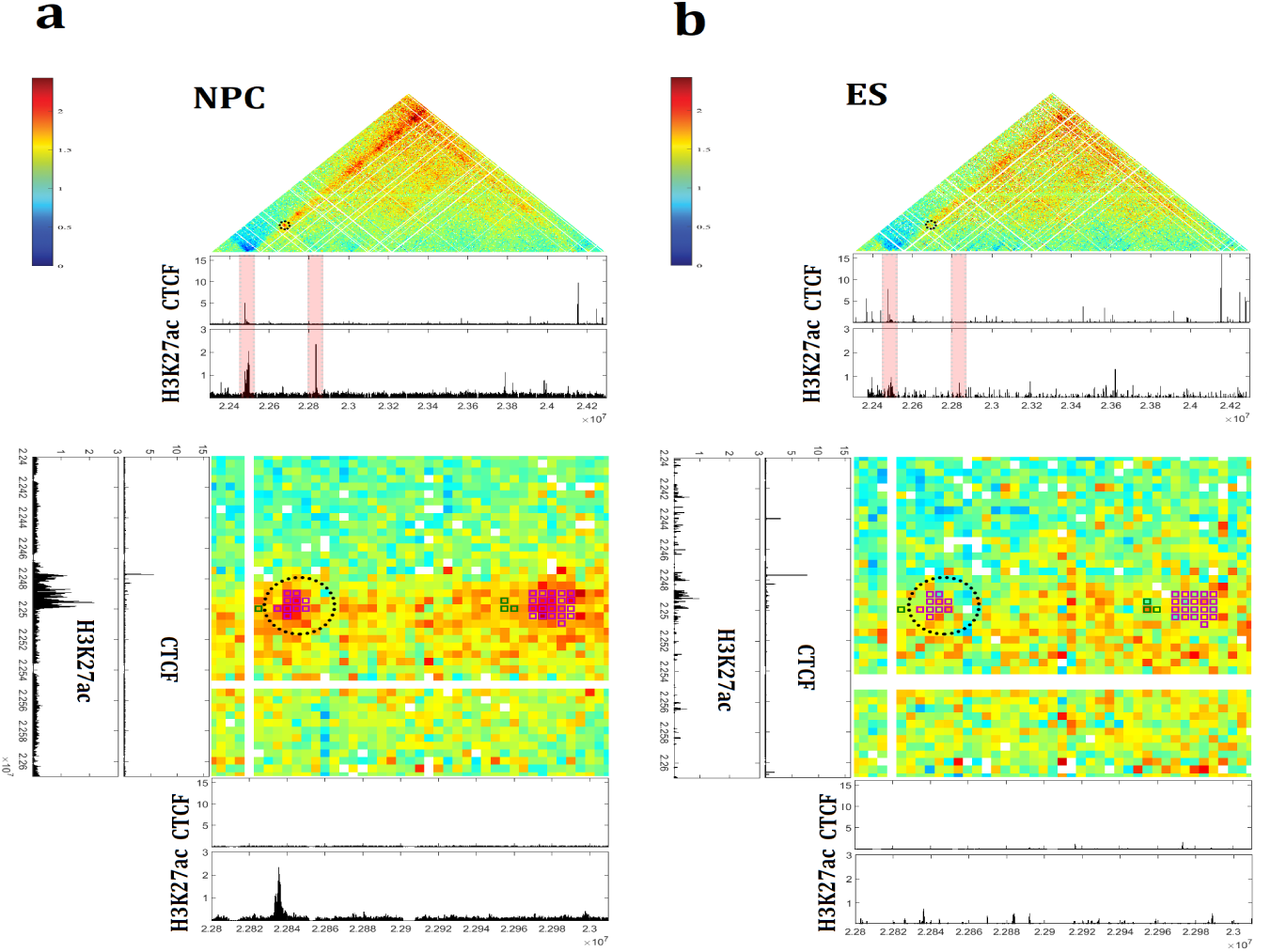
A 2-Mb region shown around Brn2 promoter of chromosome 4 of mouse neural cells (a) ES and (b) NPC. Dashed circles show the contact between the Brn2 promoter and an NPC specific enhancer. Insets show the magnified view of this contact.

## 4 Conclusion

We presented a new approach for comparative analysis of Hi-C data using a novel self-similarity measure. We showed the utility of our measure by providing solutions to two important problems in the analysis of Hi-C data, namely the problem of measuring reproducibility of Hi-C replicated experiments and the problem of finding differential chromatin interactions between two contact maps.

We showed that a simple binary comparison operation between blocks in the contact maps can be used to encode the local and global features in a manner that is robust to the data resolution and sequencing depth. This encoded information is used to build a feature vector for each contact map, which in turn allows to define a simple but effective similarity metric using the distance between their feature vectors. Experimental results showed that our self-similarity based measure outperformed two state-of-the-art methods (HiCRep and GenomeDISCO) for measuring reproducibility of replicated Hi-C experiments.

We also introduced a new method for finding differential chromatin interactions between two contact maps. SELFISH is designed based on the idea that each pairwise chromatin interaction can be represented by its neighboring interactions. Therefore, each interaction difference reveal itself as a weighted impact on the neighboring interactions. We capture this impact using a set of gradually increasing Gaussian filters. By extensively testing SELFISH on simulated and real test data we show that it outperforms the state-of-the-art DCI detection method FIND both in accuracy and efficiency.

